# Gut Microbiota and DTI Microstructural Brain Alterations in Rodents Due to Morphine Self-Administration

**DOI:** 10.1101/2024.08.15.608127

**Authors:** Kaylee Brunetti, Zicong Zhou, Samia Shuchi, Raymond Berry, Yan Zhang, Michael S. Allen, Shaohua Yang, Johnny Figueroa, Luis Colon-Perez

**Affiliations:** Department of Pharmacology and Neuroscience, University of North Texas Health Science Center, 3500 Camp Bowie Blvd, Fort Worth, TX, 76107, United States of America; Department of Microbiology, Immunology & Genetics, University of North Texas Health Science Center, 3500 Camp Bowie Blvd, Fort Worth, TX, 76107, United States of America; Center for Health Disparities and Molecular Medicine and Department of Basic Sciences, Physiology Division, Department of Basic Sciences, Loma Linda University Health School of Medicine, Loma Linda, CA, 92350, United States of America

**Keywords:** Morphine, small animal imaging, Diffusion MRI, gut, microbiota, IV self-administration

## Abstract

The opioid epidemic is an evolving health crisis in need of interventions that target all domains of maladaptive changes due to chronic use and abuse. Opioids are known for their effects on the opioid and dopaminergic systems, in addition to neurocircuitry changes that mediate changes in behavior; however, new research lines are looking at complementary changes in the brain and gut. The gut-brain axis (GBA) is a bidirectional signaling process that permits feedback between the brain and gut and is altered in subjects with opioid use disorders. In this work, we determine longitudinal, non-invasive, and in-vivo complementary changes in the brain and gut in rodents trained to self-administer morphine for two weeks using MRI and 16S rDNA analysis of fecal matter. We assess the changes occurring during both an acute phase (early in the self-administration process, after two days of self-administration) and a chronic phase (late in the self-administration process, after two weeks of self-administration), with all measurements benchmarked against baseline (naïve, non-drug state). Rats were surgically implanted with an intravenous jugular catheter for self-administration of morphine. Rats were allowed to choose between an active lever, which delivers a single infusion of morphine (0.4 mg/kg/infusion), or an inactive lever, which had no consequence upon pressing. Animals were scanned in a 7T MRI scanner three times (baseline, acute, and chronic), and before scanning, fecal matter was collected from each rat. After the last scan session, a subset of animals was euthanized, and brains were preserved for immunohistochemistry analysis. We found early changes in gut microbiota diversity and specific abundance as early as the acute phase that persisted into the chronic phase. In MRI, we identified alterations in diffusivity indices both within subjects and between groups, showing a main effect in the striatum, thalamus, and somatosensory cortex. Finally, immunohistochemistry analyses revealed increased neuroinflammatory markers in the thalamus of rats exposed to morphine. Overall, we demonstrate that morphine self-administration shapes the brain and gut microbiota. In conclusion, gut changes precede the anatomical effects observed in MRI features, with neuroinflammation emerging as a crucial link mediating communication between the gut and the brain. This highlights neuroinflammation as a potential target in addressing the impacts of opioid use.

## Introduction

Opioid use disorder (OUD) is a critical health crisis in need of novel interventions^1^. OUD is characterized by several maladaptive changes in the dopaminergic system, which is known to be of great significance to its psychopathology^2^; however, new insights provide new clues regarding the gut’s contribution to the pathology OUD. Opioid receptors are widely expressed throughout the gastrointestinal tract of rodents and humans. Activation of the μ-opioid receptor stimulates smooth muscles to reduce motility, leading to opioid-induced constipation^3–5^. Moreover, peptides in appetite regulation are expressed throughout the brain’s reward circuitry^6^, implying a synergy in the neurobiological mechanisms between food and drug consumption^7^. The gut and brain are physically connected through the vagus nerve and chemically connected through hormones, neurotransmitters, and immunological factors^8,9^ leading to bidirectional communication between the brain and gut that can influence behavior and intestinal health, which is particularly susceptible to opioid use.

Morphine is a widely used opioid for pain management with an affinity to all opioid receptors but mainly deriving its analgesic effects due to μ-opioid receptor activation^10,11^. Morphine affects the brain by inhibiting GABAergic neurons in the ventral tegmental area (VTA), reducing its inhibitory effects on dopamine (DA) neurons in the VTA, which leads to the increase of dopamine in the nucleus accumbens (NAc)^12^. On the other hand, morphine can decrease gastric motility by inhibiting emptying, increasing the sphincter tone, and blocking peristalsis^13^. In addition, morphine’s analgesia is modulated by microbiota diversity, particularly the depletion of *Bifidobacteria* and *Lactobacillaceae*^14^. Probiotic supplementation minimizes analgesic tolerance to morphine use and preserves *Bifidobacteria* and *Lactobacillaceae* communities^14^. One study showed that morphine altered several taxonomic groups in rodents’ guts, decreasing *Bacteroidates* and increasing *Flavobacterium*, *Enterococcus*, *Fusobacterium*, *Sutterella*, and *Clostridium* ^15^. Knocking down gut microbiota with antibiotics alters the NAc and prefrontal cortex transcriptional responses to morphine and decreases the formation of morphine conditioned place preference and locomotor sensitization^16^. Overall, morphine’s action induces alterations in the brain and gut that modulate behavior and increase the risk for substance use disorders.

Morphine’s simultaneous effects in the periphery and the central nervous system make it necessary to determine simultaneous alterations in the brain and gut. One relevant technique to study the brain is Magnetic Resonance Imaging (MRI). MRI is a well-established approach to studying the neurobiology of neuropsychiatric disorders like addiction and SUDs^17^. MRI is a non-invasive and translational approach that enables researchers and clinicians to gain complementary insights into human clinical trials and essential neuroscience experimentation. In addition, sequencing technologies that allow the microbiome to be investigated *in vivo* and non-invasively, such as 16S rDNA sequencing of fecal matter, can yield instrumental insights for a comprehensive understanding of the relationship between changes in the brain and the gut^18^. The gastrointestinal tract contains 10^14^ microbiota, and the total genes it comprises are 100 times that of the human genome^19^; therefore, determining how the gut and its microbiome are affected by opioids alongside markers of brain health is critical to advancing our understanding of SUDs. The flexibility of MRI and the readily available access to fecal analysis allows us to obtain longitudinal data on brain and gut health simultaneously.

Given the ongoing evolution of gut health studies and the role of microbiome diversity in SUDs, it is crucial to assess further the direct effects of opioids on the gut and brain. Neuroimaging is a critical tool to determine the concurrent neurobiological alterations due to opioid use that can be related to simultaneous changes in gut composition associated with the use of opioids; however, neuroimaging and gut diversity have *not* been explored in the context of SUD. Given the many intersections between the gut and brain in developing SUDs, neuroimaging could be a critical approach combined with gut health assays to gain new insights about SUDs. This article utilizes animals’ self-administering morphine to determine concurrent alteration in gut microbiome and brain features measured with dMRI. In a small subset of animals, we identify potential evidence of neuroinflammation in the brain that could mediate changes in MRI. This work presents links for the hypothesis that the brain and gut continuously communicate with one another when consuming drugs of abuse. Further expanding our knowledge on GBA by utilizing neuroimaging in animal models of volitional substance use could provide new insights into the modification of the gut and brain due to prolonged morphine use.

## Methods

### Animals

Adult male (n = 13) and female (n = 13) Sprague Dawley rats were purchased from Charles River Laboratories (Raleigh, North Carolina). Upon arrival, they were housed in pairs on a reverse light/dark cycle in controlled rooms for temperature and humidity. Food and water were available ad libitum. The Department of Laboratory Animal Medicine at the University of North Texas Health and Science Center provided care for rats in this study, and the Institutional Animal Care and Use Committee at the University of North Texas Health and Science Center approved all experiments.

### Surgery and Catheter Implantation

Rats were quarantined for roughly a week before undergoing catheterization surgery. Rats were anesthetized with isoflurane (5% induction, 2% maintenance) and administered subcutaneously Meloxicam (1 mg/kg). The right jugular vein was ligated with a catheter and secured in place using aseptic surgical techniques, ^20^. The jugular catheters were constructed in-house with MRI-compatible cannulas (P1 Technologies, Roanoke, VA). The tubing side of the catheter was passed subcutaneously over the right shoulder and through a small incision in the skin over the scapulae. The cannula was secured to the back of the rats using a surgical mesh to hold them in place. The skin around the backport was then sutured, and a protective aluminum cap was placed on the port (McMaster Carr). Rats were given 5 d to recover from surgery, after which they were shaped for morphine or sucrose pellet self-administration. Catheters were checked for patency by observing a rapid and transient loss of muscle tone following an infusion of 0.1 ml of propofol.

### Intravenous Self-Administration of Morphine

Self-administration operant behavior was performed in eight identical rat behavior chambers (MedAssociates, Fairfax, VT, USA). The chambers were extra tall with a modified top enclosure housed in a PVC sound-attenuating cubicle. The chambers were equipped with a house light, two stimulus lights, two retractable levers, a pellet dispenser with a trough and head entry detection, a variable speed syringe pump, an 8in/8out SmartCtrl package, and a drug delivery arm. The syringes were connected to the catheter in the rat’s back through PE50 tubing that reached the back-mounted venous access port. During self-administration sessions, only one of the two levers resulted in a successful infusion of morphine (active lever), and the other had no consequence (inactive lever). The Drug Supply Program of the National Institute on Drug Abuse supplied the morphine used in this study. Morphine hydrochloride was dissolved in 0.9% sterile saline (0.4 mg/kg/infusion). All experiments were performed during the active/dark cycle phase. Before drug self-administration, the animals had four training phases consisting of 1) habituation with sucrose pellets in the food dispenser, 2) magazine training, 3) lever pressing for sucrose pellets under a fixed ratio (FR)1-L, and 4) lever training and FR1-R lever training. Once the training phases were completed, the animals started their two-hour sessions of drug self-administration (morphine, 0.4mg/kg). During drug self-administration, both levers were extended during the duration of the two-hour sessions, but FR1-R was programmed to be the lever associated with drug administration. Once the animals pressed FR1-R, a cue light would illuminate, and animals would receive 0.01 mL of morphine per lever press. Before rats began administering drugs, their catheters were flushed with 0.1 mL of saline to confirm that the catheters were working. After each behavioral session, the catheters were flushed with 0.2ml of Cefazolin to prevent infection and 0.1ml of Heparin. All behavioral experiments were performed in MED-PC. Rats from either group had to meet a criterion of drug or pellet seeking from day 7 to day 14 but pressed at least seven times for morphine/pellet to be considered for behavior and MRI analysis.

### Fecal collection

Fresh fecal samples were collected from awake animals at three different time points throughout this experiment. Fecal collection occurred at (1) baseline, seven days after catheter implantation; (2) acute, 24 hours after the second day of self-administration or the third day experimentation (no IVSA on this day); and (3) chronic, 24 hours after completion of the 12 days of self-administration. Prelabeled collection tubes were prepared with the date, rat number, and trial number before fecal collection. A clean cage was obtained from the vivarium and sanitized with 99% isopropyl alcohol between each rat. New gloves were worn for each fecal collection and sprayed with 99% isopropyl alcohol; face masks were worn to avoid contaminating the fecal samples. Fecal samples were collected and immediately stored at -80°C until DNA extraction.

### 16S rDNA Sequencing

Microbial sequencing was performed as previously described in.^21,22^ Microbial DNA was extracted from 50∼100 mg fecal material using the Qiagen DNeasy PowerSoil Pro Kit and the automated QIAcube Connect robot (Qiagen, Carlsbad CA) following the manufacturer’s instructions. We used universal bacterial primers to target the V4 hypervariable region and amplify the 16S rRNA gene ^23,24^. The PCR reaction contained: DNA template (10 ∼ 100ng), 0.5 μL of each primer (10 μM), 2.5 μl 10X AccuPrime PCR Buffer II, 2.5 mL BSA (1.6 mg/mL), 1.5 μl Mg (50 mM), 0.1 μL AccuPrime Taq High Fidelity (5U/μL), and PCR grade water to a final volume of 25 μL. PCR amplification was performed in duplicate for each sample and carried out as follows: heated lid 94°C for 2mins, 25 cycles of 94°C for 30s, 52°C for 30s, 68°C for 40s, then 68°C for 5 mins and held at 4°C. The PCR products were confirmed with gel electrophoresis (1.5% agarose), duplicate reactions were combined, and successful products were cleaned using AMPure XP magnetic bead-based purification. We used Illumina Nextera XT Index Kit v2 (Illumina, San Diego, CA), following the manufacturer’s instructions, to index the clean PCR products and repurified them with AMPure XP magnetic beads (Beckman Coulter, Chaska, MN). Each 50 μL index PCR reaction contained 5 μL 10X AccuPrime PCR Buffer II, 5 μL Nextera XT indexing primers 1, 5 μL Nextera XT indexing primers 2, 0.2 μL AccuPrime Taq High Fidelity (5U/μL), 5 μL purified DNA, and PCR grade water. The PCR recipe was as follows: heated lid 94°C for 3mins, 8 cycles of 94°C for 30s, 55°C for 30s, 68°C for 30s, then 68°C for 5 mins and held at 4°C. We used the Qubit dsDNA HS Assay Kit (Invitrogen, Carlsbad, CA) to quantify PCR products and pool them in equal amounts. The pooled sample was then denatured, diluted, loaded, and sequenced using a Miseq Reagent V2 (500 cycles) kit following the manufacturer’s instructions.

#### Bioinformatics Analysis

The generated DNA sequences were analyzed using the mothur MiSeq SOP pipeline^25^. We assembled paired-end sequences and removed short (< 100bp) and low-quality sequences (homopolymers > 8) from the dataset. We used the SILVA database to align sequences. The unaligned sequences and gaps were removed. Redundant sequences were reduced using the unique.seqs command, a precluster (diffs=2) algorithm, and chimeras were removed after identification using UCHIME^26^. We used the Ribosomal Database Project (RDP) classifier with a minimum of 80% confidence to classify taxons ^27^ and the up-to-date curated EzBiocloud database as a reference. We removed sequences classified as mitochondria, chloroplast, archaea, and eukaryote from the dataset.

Microbial diversity (Shannon diversity and evenness), richness (Chao1), and abundance coverage-based estimator (ACE) were calculated based on Amplicon sequence variants (ASVs) ^28–30^. Microbial communities between morphine and pellets groups were compared and visualized using UniFrac distances and principal coordinate analysis (PCoA)^31^. The Molecular Variance analysis (AMOVA) was performed to assess the variability among and within different groups^32^. Diversity estimators, UniFrac, Principal Coordinate Analysis (PCoA), and Analysis of Molecular Variance (AMOVA) analyses were performed using mothur.

### Diffusion Magnetic Resonance Imaging

Rats (n = 20) were scanned in a cryogen-free MRI magnet (MRS*DRYMAG7017, MR Solutions, UK) capable of changing between 3T and 7T fields with a 17-cm bore magnet equipped with a 600 mT/m RF gradient. All studies were performed at the 7T field strength. A quadrature transmit/receive radio frequency (RF) coil was used for B1 field excitation and RF signal detection, designed explicitly for rat head imaging. All rats underwent three imaging sessions (Fig 1A). Session one of imaging occurred on day 0, before the administration of drugs or behavioral training. Session two of imaging was on day three of the experiment, after allowing the animals to administer drugs for two days. Session three was on day fifteen of the experiment after the animals finished 14 days of drug self-administration. The animals were in a controlled room with a reverse light/dark cycle while waiting in all three imaging sessions. The animals were anesthetized with isoflurane (5% induction, ∼1.5% maintenance) and then transferred to the scanner, where temperature and breathing rate were monitored throughout the scan (SA Instruments). A warm air-heated bed maintained the rat’s body temperature at 37– 38°C. Diffusion-weighted images were collected using a two-shot spin-echo echo planar imaging (EPI) sequence with the following parameters: 2 diffusion weighting shells of 18 directions with Jones arrangement^33^ at b = 500, and 60 Jones with b = 900 s/mm^2^, and 4 b=0 images, echo time (TE) = 25 ms; repetition time (TR) = 5.0 s; 32 × 32 mm in plane; 16 slices with 1.3 mm thickness per slice; data matrix = 100 × 94; phase encode direction IS. We also obtain a single b = 0 blip-down image with phase encoding in the SI direction. Anatomic scans for image overlay and reference-to-atlas registration were collected using a FLASH 3D sequence (TI = 2200 ms; TE = 4.9 ms; TR = 25 ms; number of averages = 3; data matrix = 134 x 128 × 100 and spatial resolution = 0.27 x 0.25 x 0.3 mm^3^).

**Figure 1.**
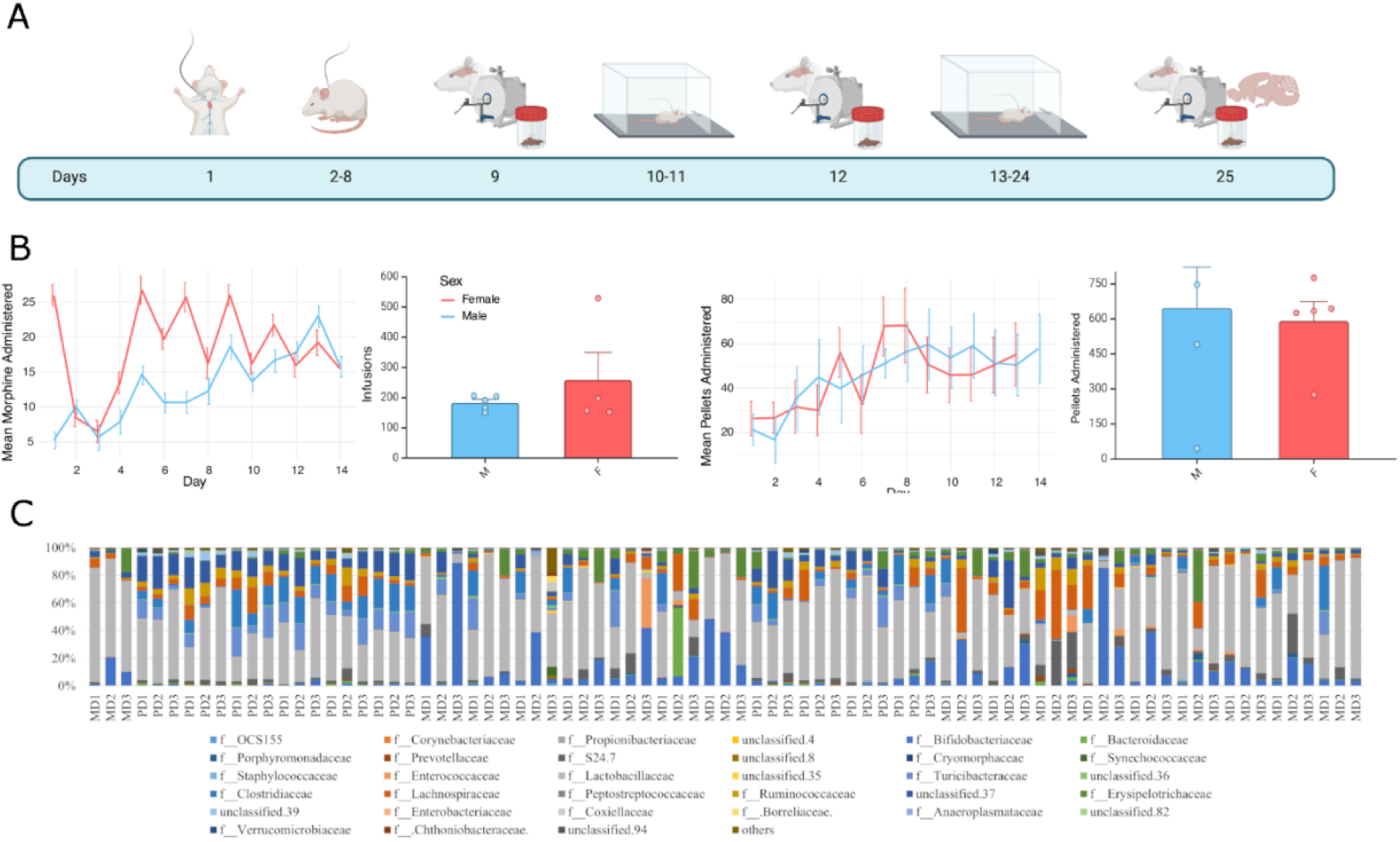
Morphine self-administration and microbiome composition. A) Experimental timeline: rats undergo catheter implantation (day 1) and a recovery period (days 2-8) before experimental procedures. All rats were probed for baseline gut and brain conditions with fecal collection and MRI (day 9). Then rats undergo acute self-administration for 2 days before a second probing session of gut and brain (day 12), then rats continue to a chronic phase where they undergo SA for an additional twelve days before the final and chronic probing sessions that include feces collection, MRI, and brain tissue. B) Self-administration: a daily average of infusions or pellet delivery and total number of infusions during the 14 days or pellets delivered. All operant responses showed comparable results for all animals of either sex. C) Normalized community composition of all animals at all time points at the family level. M = morphine, P = pellets, D1 = day 1, D2 = day 2, and D3 = day 3.

### DTI image processing and analysis

Brain masks were generated using high-resolution anatomic scans, and RBM was manually edited to maintain only brain tissue using ITK-SNAP. The cropped brain structural image was aligned with the B0 image of the DWI scan to obtain a brain mask in DWI space using the FMRIB Software Library linear registration program FLIRT. Diffusion data was stripped of the skull and corrected for field inhomogeneities (FSL’s topup). The DWI image was registered to the SIGMA rat brain template using ANTs (AntsQuickSyn) to obtain segmentation and perform voxel-based analysis and seed-based analysis. We generated DTI indices (FSL’s dtifit): fractional anisotropy (FA), mean diffusivity (MD), radial diffusivity (RD), and axial diffusivity (AD) in native space. The voxel-based analysis was completed in an in-house template of FAs including the baseline data from both groups, using ANTs function antsMultivariateTemplateConstruction2.sh. Then, we proceeded to register all FAs of each sample in both groups and in all days: baseline, acute, and chronic, onto the in-house template-FA, using ANTs function antsRegistrationSyn.sh. Then, we applied the in-house template registration onto MD, AD and RD, so that all DTI maps are warped in the in-house template image space using ANTs function antsApplyTranform.sh. Then we t-threshold = 2.11 according to the degree of freedom = 10 + 10 - 2 in these tests. Once their t-scores are computed, the corresponding p-values are computed. FA, AD, RD, and MD map voxel-level t-tests (alpha = 0.05) were completed using Python’s Numpy and Scipy packages with voxel-wise t-tests (alpha = 0.05) followed by Family-Wise Error (FWE) rate correction using the Benjamini-Hochberg False Discovery Rate into q-values < 0.05. Finally, we registered the SIGMA labels onto the in-house template and back to the native space of each individual samples to perform seed-based analysis. Seed-based analysis was analyzed using Welch Two Sample t-tests on the mean FA, AD, and MD maps in R with the t-test function and shown with a box plot using the ggplot function.

### Tissue Collection and Immunohistochemistry

The animals were anesthetized with isoflurane and decapitated. Brains were rapidly removed, fixed in 4% formalin, and further processed to be embedded in paraffin. The immunofluorescence protocol used in this study was adapted from Liu et al^34^. Briefly, brain sections were obtained (10-μm thick) using a microtome. Deparaffinization was done for antigen retrieval, and tissue was placed in a citric acid buffer in a boiling water bath. Sections were washed with 0.2% triton x-100 in PBS and blocked using 10% normal goat serum in superblock for an hour and thirty minutes at room temperature. Brain sections were then stained with primary antibodies for GFAP (Santa Cruz, 1:200), Iba1 (Cell Signaling, 1:200), and DAPI (Cell Signaling, 1:200) overnight at 4 °C, followed by species-specific secondary antibodies. Microscopic images were acquired with a Keyence BZ microscope and analyzed with FIJI^35^.

### Immunofluorescence analysis

In our study, we used the FIJI AnalyzeSkeleton plugin to calculate the area covered, branch size, and branch length, to characterize cell morphology following morphine IVSA. Results were analyzed with a Mann-Whitney U Test^36^.

## Results

### Morphine self-administration

The animals self-administered morphine on average 13.0 ± 1.8 daily infusions for males and 18.4 ± 12.9 daily infusions for females (Fig 1B). The main discrepancy in the high variability of female drug intake was one rat that self-administered an average of 38 daily infusions. Overall, females self-administered on average 257.8 ± 181.3 infusions, and males self-administered 182.4± 24.6 (p = 0.468, Fig 1B). The control group also successfully pressed levers for sucrose pellets, with daily averages of 46.1 ± 27.9 pellets for males and 45.3 ± 14.4 pellets for females (Fig 1B). Overall, females consumed 645.2 ± 391.2 pellets over the 14 sessions, and males 589.2 ± 186.6 (p = 0.783, Fig 1B). There was no significant difference in drug consumption between sexes; hence, going forward, we aggregate all results in terms of morphine vs the control group.

### Gut Microbiome

Bacterial DNA analysis of fecal matter showed microbial diversity changes in rats self-administering morphine. There were significant differences in alpha diversity measures across days among the pellet and morphine groups. Chao1 showed a progressive reduction in diversity with a significant difference in abundance between baseline and the chronic stage in the morphine group (p=0.024) but not for the sucrose pellet (Fig 2A). Chao1 also displays differences between groups at the acute stage (p=0.048) and chronic stage (p=0.031, Fig 2A). The Shannon index showed a significant difference in species diversity between groups at the acute stage only (p=0.010, Fig 2B). The Evenness index showed no significant changes across days or treatment (Fig 2C). In terms of beta diversity, there were noticeable differences in the principal coordinates analysis (PCoA) across days and whether they administered morphine or sugar pellets with the baseline and of the morphine group aligning with all stages of the sugar pellets between the 1^st^ and 2^nd^ axis (Fig 2D) and 1^st^ and 3^rd^ axis (Fig 2E).

**Figure 2.**
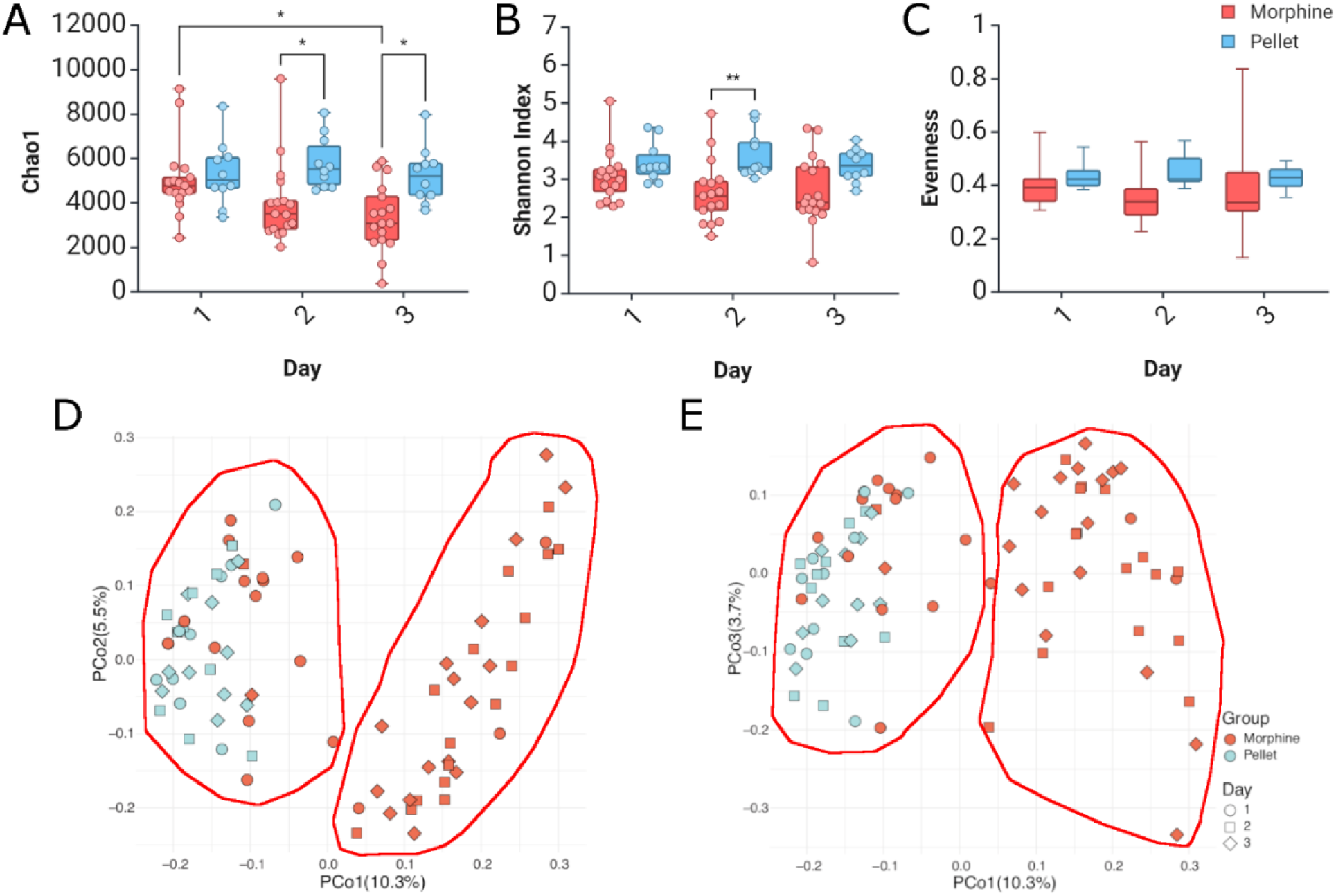
Diversity changes and clustering of microbes due to morphine SA. A) Chao1, B) Shannon index, and C) evenness results reflecting relative abundances for each timepoint (i.e., baseline = Day 1, acute = Day 2, and chronic = Day 3) and group (i.e., morphine vs sugar pellet). Principal coordinate analysis (PCoA) based on unweighted unifrac. * p < 0.05

Diversity alteration reflected changes at the different taxonomic levels. We found that the phylum *Firmicutes* showed significant changes in percent abundance within the morphine group between baseline and the acute phase (p=0.009) and between groups at the acute phase (p=0.003) and chronic stage (p=0.026, Fig 3A). The phylum Bacteriodetes did not show any differences at any stage or between groups (Fig 3B). The family Bifidobacteriaceae showed significant changes in abundance between groups at the acute stage (p=0.030, Fig 3C). The family Erysipelotrichaceae showed substantial changes in abundance within the morphine group between baseline and chronic stages (p=0.039, Fig 3D). The families Ruminococcaceae (Fig 3E) and Lactobacillaceae (Fig 3F) did not show any significant differences in abundance across treatments or days. The differences between operational taxonomic units at the family levels show that the alterations are more prominent for the morphine group than controls (brighter colors of red and green in Fig 4A-C). The between-group differences in OTUs at the family level between groups and stages show that differences become more prominent after the baseline when animals self-administer morphine (Fig 4D).

**Figure 3.**
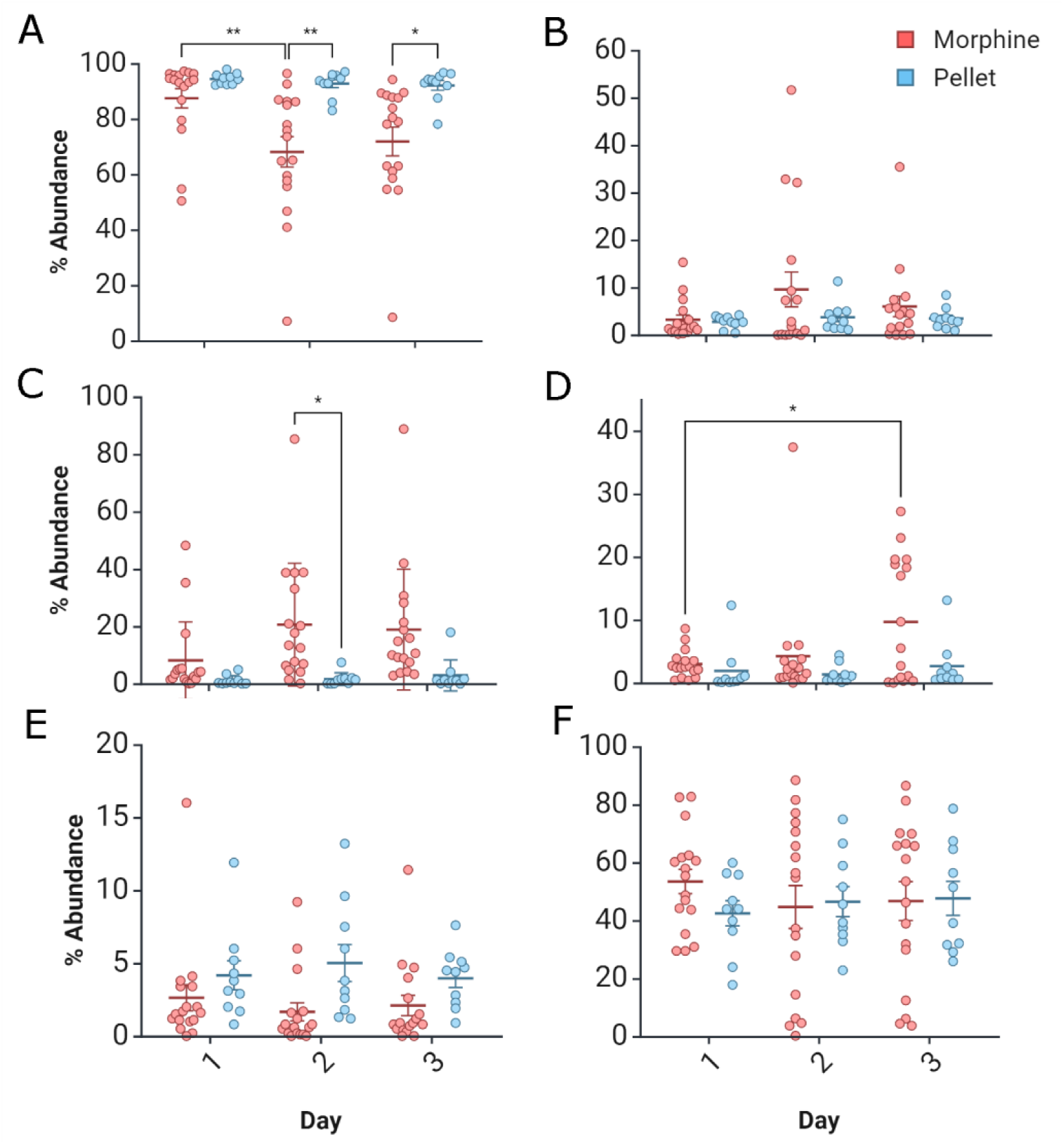
Phylum and family relative abundances for each timepoint (i.e., baseline = Day 1, acute = Day 2, and chronic = Day 3) and group (i.e., morphine vs sugar pellet). Phylum level changes for A) Firmicutes and B) Bacteriodetes. Family level changes for C) Bifidobacteriaceae, D) Erysipelotrichaceae, E) Ruminococacea, and F) Lactobacillaceae. * p < 0.05 decrease in Firmicutes and increase in Bifidobacteriaceae, and Erysipelotrichaceae.

**Figure 4.**
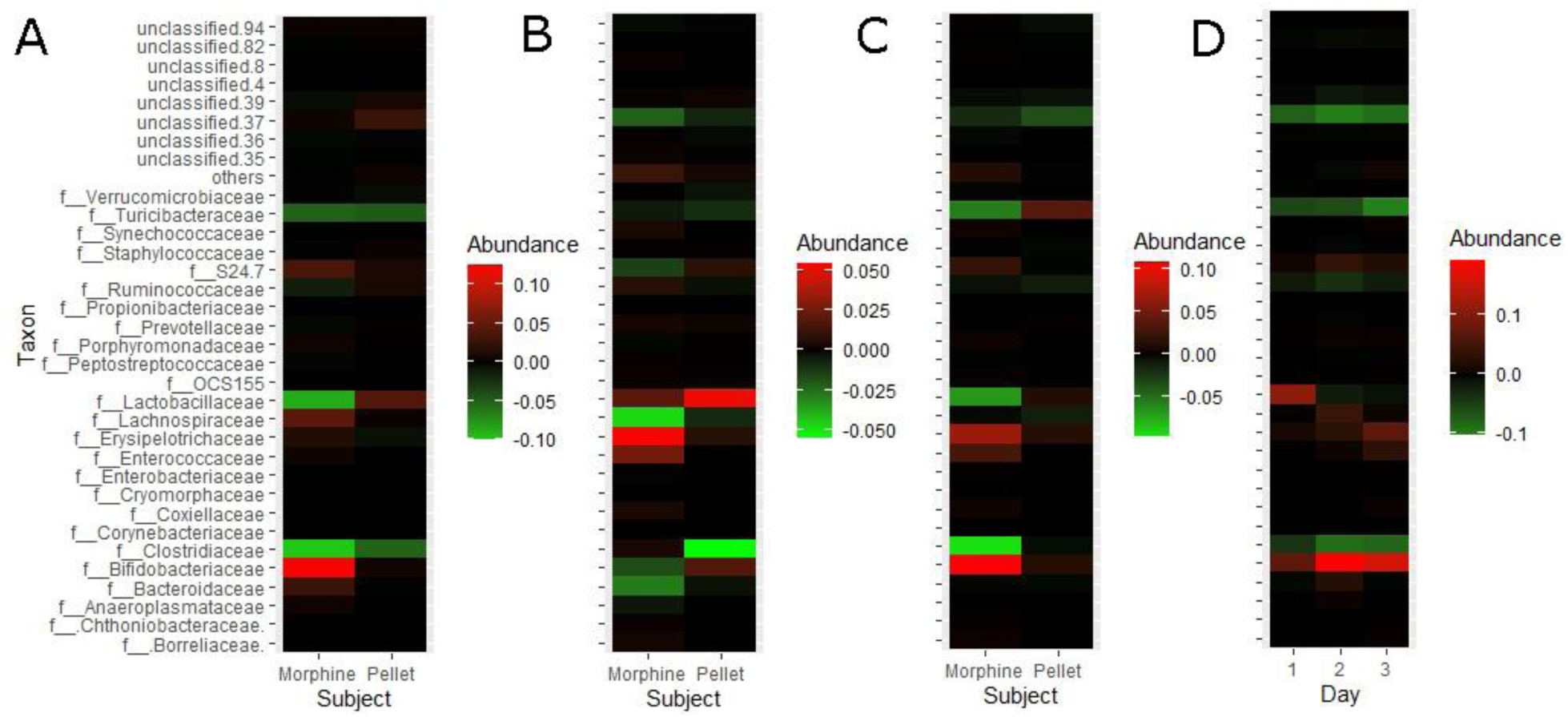
Differences in relative abundances of OTUs at the family level. A) Acute minus baseline (red acute higher concentration and green baseline higher concentration), B) Chronic minus baseline (red chronic higher concentration and green baseline higher concentration), and C) Chronic minus acute (red chronic higher concentration and green acute higher concentration). D) comparison of group differences (morphine minus saline, red morphine higher, and green pellet higher) for each day. Each timepoint (i.e., baseline =Day 1, acute = Day 2, and chronic = Day 3) and group (i.e., morphine vs sugar pellet).

The phylum and family alteration reflect changes also observed at the genus level. The genus *Clostridium* showed significant changes in abundance between groups at the acute stage (p=0.004) and chronic state (p=0.049, Fig 5A). The genus *Ruminococcus.1* showed a significant difference between groups at the acute stage (p < 0.001) and chronic (p=0.002, Fig 5B). The genus *Bifidobacterium* showed alterations between groups at the acute stage (p = 0.030). The genus *Turicibacter* showed significant changes in percent abundance between groups at the chronic stage (p= 0.001). The genus *Allobaculum* abundance did not show any significant differences in abundance across treatments or days.

**Figure 5.**
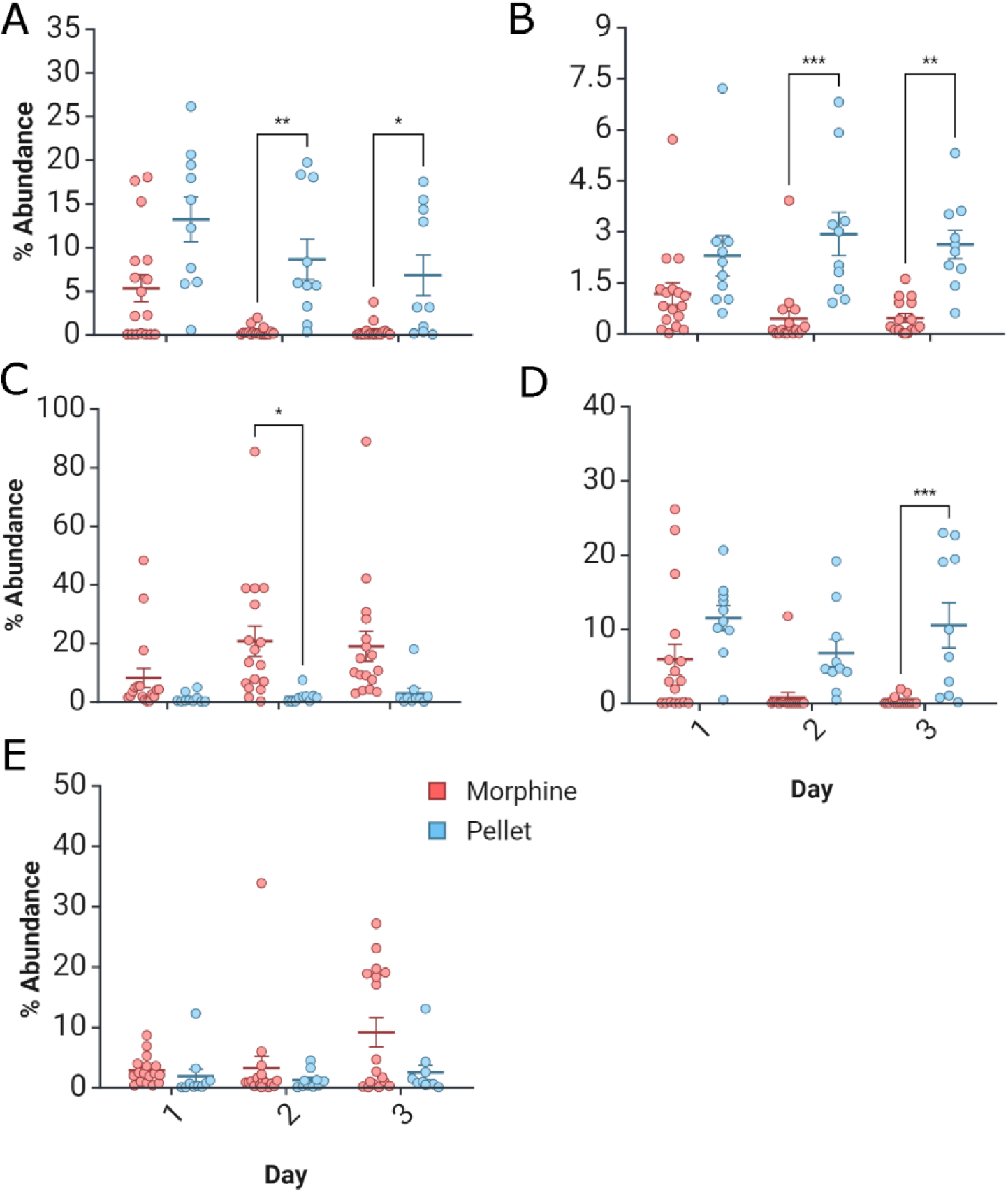
Genus relative abundances for each timepoint (i.e., baseline = Day 1, acute = Day 2, and chronic = Day 3) and group (i.e., morphine vs sugar pellet). A) *Clostridium*, B) *Ruminococcus 1*, C) *Bifidobacterium*, D) *Turicibacter*, and E) *Allobaculum*. * p < 0.05

### MRI

Voxel-based analysis of DTI indices showed longitudinal changes within the morphine and pellet self-administering groups along between groups comparison at the chronic stage. The FA changes were observed for frontal and striatal regions (Fig 6). The striatum display a robust within effect between acute and chronic stage, but not between baseline and acute. Between subject shows a stark contrast in the chronic stage replicating within subjects’ findings. Other areas showing differences are the septal region, the primary cingular cortex (PCC), and the corpus callosum (CC). The pellet group did not show large differences from the morphine group, and the early stages of the between-subjects test also did not show major alterations between groups.

**Figure 6.**
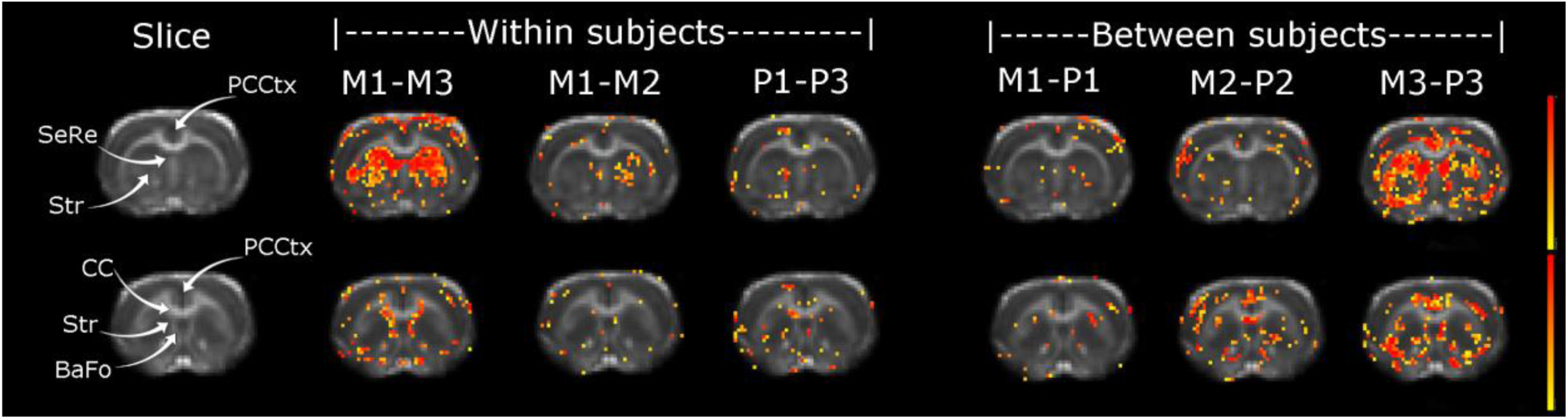
Voxel based differences of FA maps. The morphine treatment induced chronic change in FA, confirmed by a between groups change in striatum. The acute and pellet state did not induce extensive alterations. PCCtx = primary cingular cortex, SeRe = septal region, Str = Striatum, CC = corpus callosum, BaFo = basal forebrain region, M1 = Morphine group baseline, M2 = Morphine acute, M3 = Morphine chronic, P1 = pellet group baseline, P2 = pellet group acute, P3 = pellet group chronic.

The MD changes resemble alterations observed for FA, with major differences within subjects but not at the between-subject chronic stage (Fig 7). The lack of between subjects’ alterations for the pellet group is further explained as the pellets within subjects’ alterations between baseline and chronic are prominent for the similar areas of the morphine group. The MD differences are again prominent for the striatum, and PCC for the FA. In addition, we observe the thalamus, basal forebrain region, primary motor cortex (PMC), fimbria, retrosplenial cortex, and somatosensory cortices (SMT, dysgranular, hindlimb, and barrel field). The main differences in MD are widespread at the comparison within subjects for both groups with morphine inducing larger effects than sucrose pellets. The within subject test for the acute stage did not show changes implying an effect of chronic consumption of morphine or sucrose, with the thalamus exclusively showing differences for morphine.

**Figure 7.**
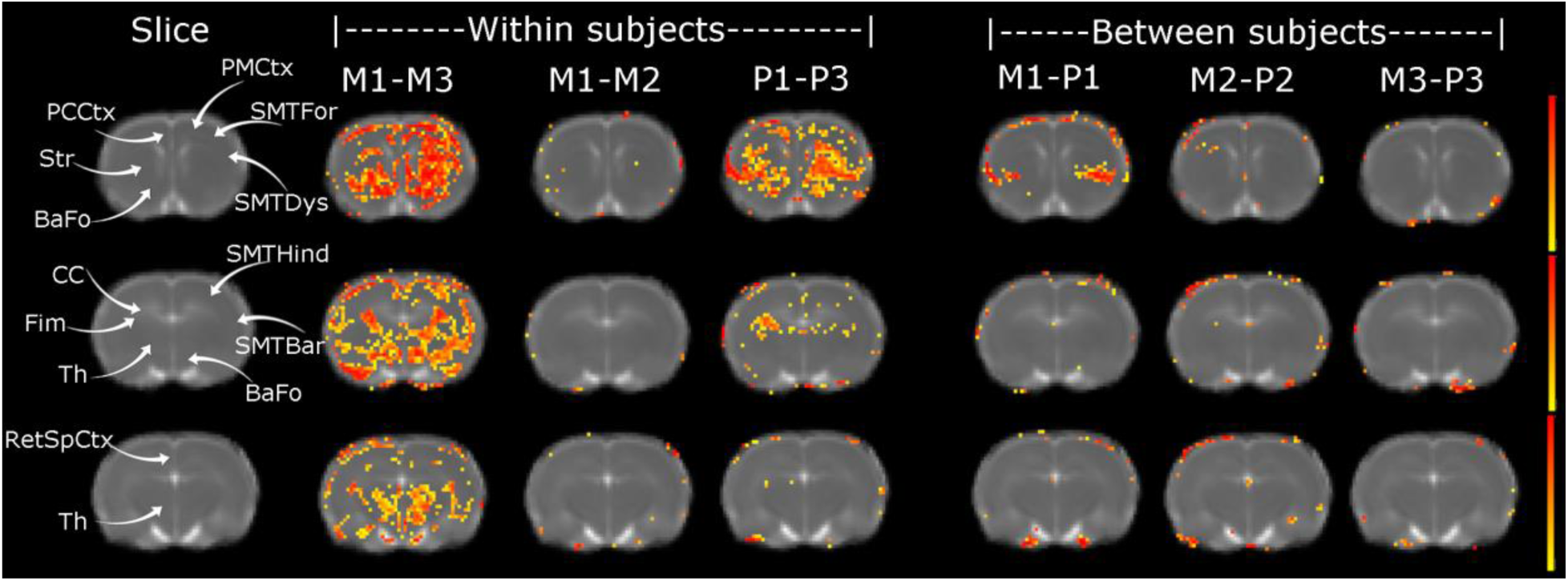
Voxel based differences of MD maps. MD longitudinal changes were most extensive between chronic and baseline groups in the morphine and pellet treatment comparisons, but morphine induced more widespread changes than pellets. MD differences did not show extensive changes in the between subject’s tests. PCCtx = primary cingular cortex, PMCtx = primary motor cortex, SMTFor = somatosensory cortex forelimb, SMTDys = somatosensory cortex dysgranular, SMTBar = somatosensory cortex barrel field, SMTHind = somatosensory cortex hindlimb, SeRe = septal region, Str = Striatum, CC = corpus callosum, Fim = fimbria, Th = thalamus, RetSpCtx = retrosplenial cortex, BaFo = basal forebrain region, M1 = Morphine group baseline, M2 = Morphine acute, M3 = Morphine chronic, P1 = pellet group baseline, P2 = pellet group acute, P3 = pellet group chronic.

**Figure 8.**
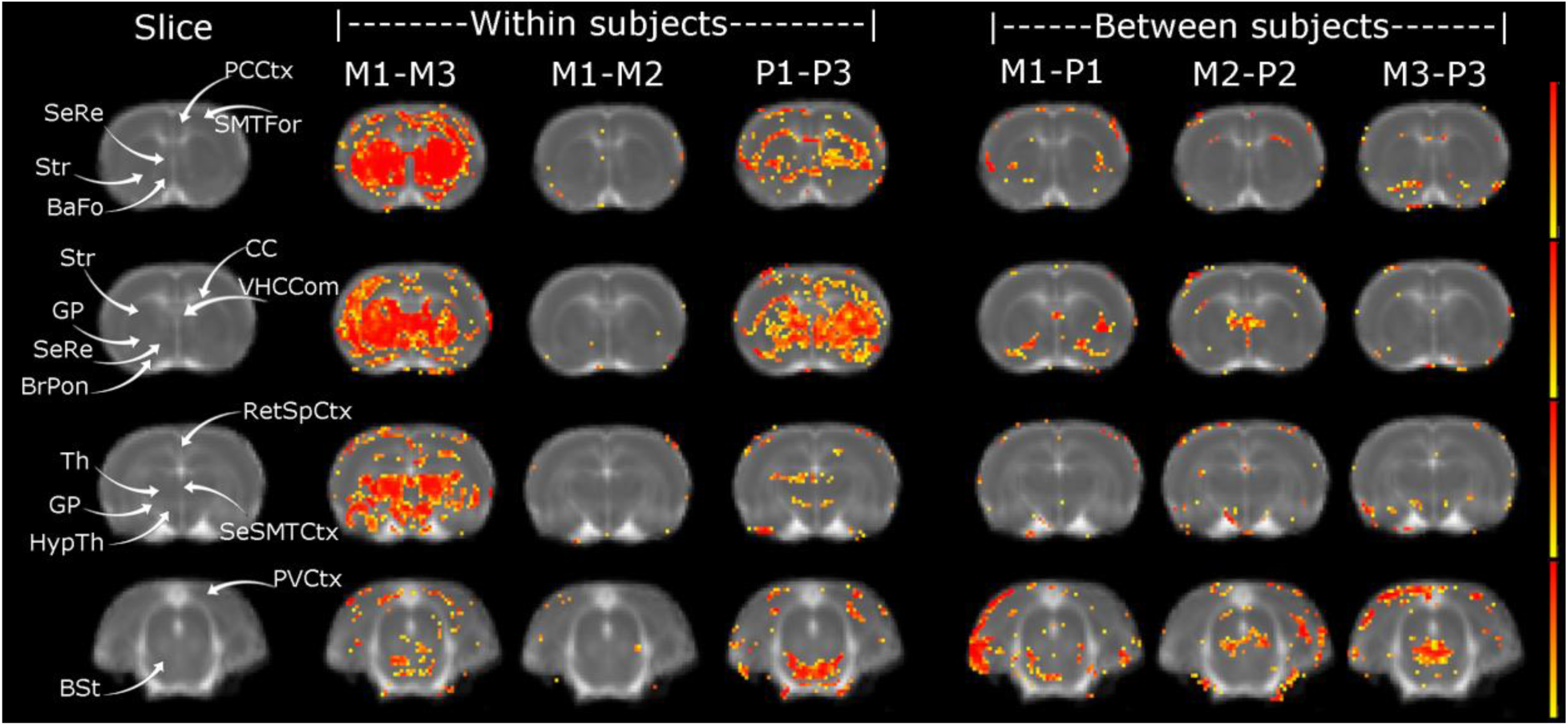
Voxel based differences of AD maps. AD longitudinal changes were most extensive between chronic and baseline groups in the morphine and pellet treatment comparisons, but morphine induced more widespread changes than pellets. AD differences showed changes in the between subject’s tests mainly in the brain stem. PCCtx = primary cingular cortex, SMTFor = somatosensory cortex forelimb, SeRe = septal region, Str = Striatum, VHCCom= ventral hippocampal commissure, BrPon = brachium pontis, CC = corpus callosum, SeSMTCtx = secondary somatosensory cortex, Th = thalamus, HypTh = hypothalamus, GP = globus pallidum, RetSpCtx = retrosplenial cortex, BaFo = basal forebrain region, BSt = brain stem, PVCtx = primary visual cortex. M1 = Morphine group baseline, M2 = Morphine acute, M3 = Morphine chronic, P1 = pellet group baseline, P2 = pellet group acute, P3 = pellet group chronic.

**Figure 9.**
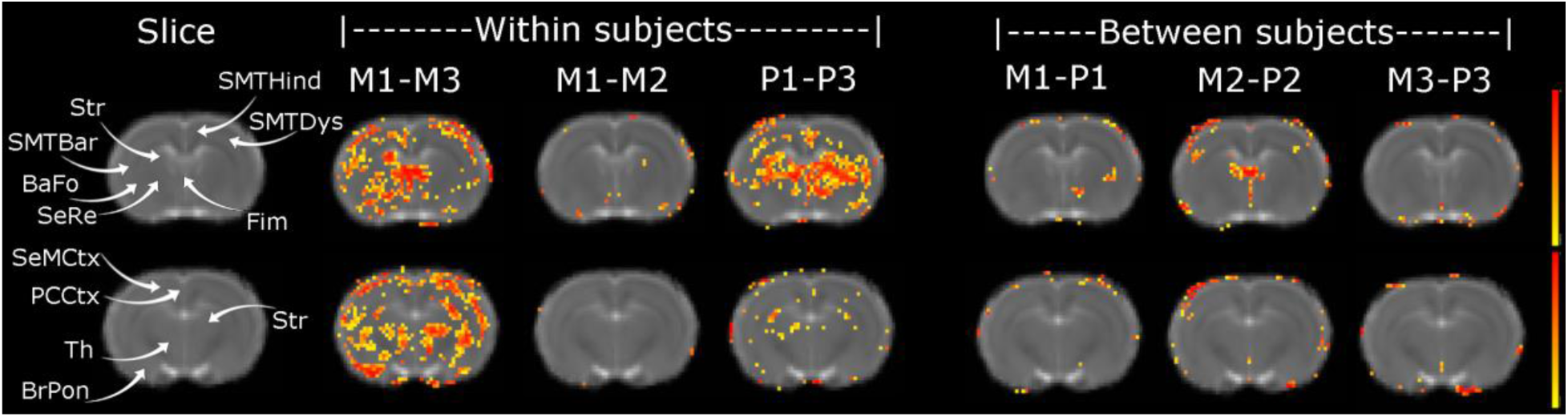
Voxel based differences of RD maps. RD longitudinal changes were most extensive between chronic and baseline groups in the morphine and pellet treatment comparisons, but morphine induced more widespread changes than pellets. RD differences did not show extensive changes in the between subject’s tests. PCCtx = primary cingular cortex, SMTDys = somatosensory cortex dysgranular, SMTBar = somatosensory cortex barrel field, SMTHind = somatosensory cortex hindlimb, SeRe = septal region, Str = Striatum, BrPon = brachium pontis, Fim = fimbria, SeMCtx = secondary motor cortex, Th = thalamus, BaFo = basal forebrain region. M1 = Morphine group baseline, M2 = Morphine acute, M3 = Morphine chronic, P1 = pellet group baseline, P2 = pellet group acute, P3 = pellet group chronic.

The AD changes are the most prominent but maintain anatomical correspondence with FA and MD. In addition to frontal and striatal regions, we identified alteration in AD for the brain stem (locations where ventral tegmental area, dorsal raphe, pontine nucleus, and ventral periaqueductal gray areas). As with the prior test, we again observe major differences in the within-subject test but less between subjects. We observe striatum and PCC, in addition to septal region, basal forebrain region, globus pallidum, brachius pontis, thalamus, hypothalamus, SMT’s forelimb areas, ventral hippocampal commissure, CC, retrosplenial cortex, secondary SMT, and primary visual cortex. The within-subject test for the acute stage did not show changes implying an effect of chronic consumption of morphine or sucrose as in the MD test.

The RD resembles changes discussed for the MD albeit less extensive. The within-subject test for the acute stage did not show changes implying an effect of chronic consumption of morphine or sucrose, and there were no significant changes between subjects. Areas displaying alterations in the RD are the striatum, SMTs hindlimb, dysgranular, and barrel field areas, basal forebrain, septal region, brachius pontis, secondary motor cortex, and thalamus.

### Immunohistochemistry

We determined markers of neuroinflammation in the thalamus and found significant alterations in rats self-administering morphine (Fig. 10A-D). The morphine self-administration group displayed reduced area, and branching, while an increase in branch lengths (Fig 10G-I). No differences in area were seen for nuclear or astrocytes (Fig. 10E-F).

**Figure 10.**
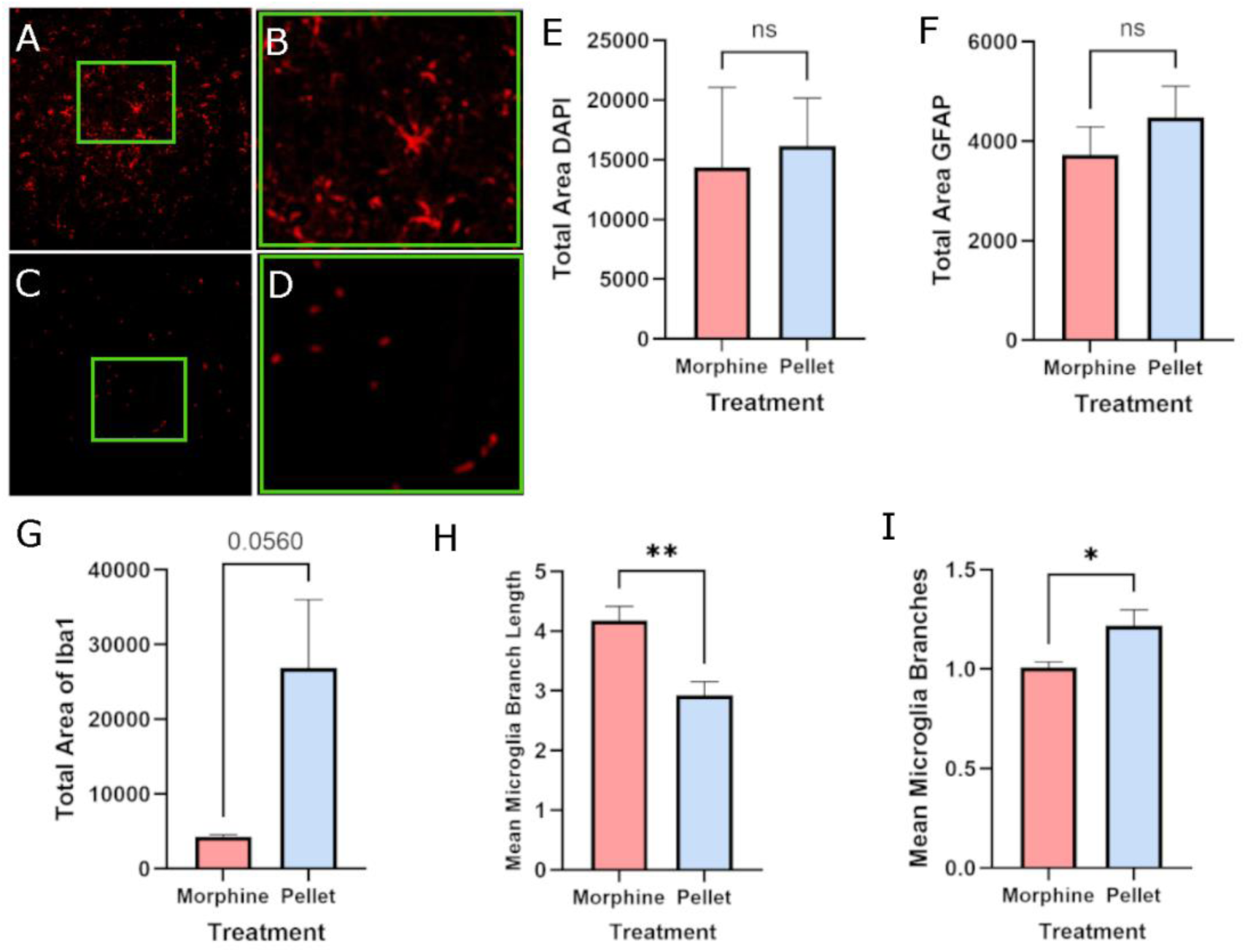
Thalamus immunohistochemistry results. A) Pellet control thalamus Iba1 staining. B) Enlarged Iba1 staining for green square in A. C) Morphine group thalamus Iba1 staining. D) Enlarged Iba1 staining for green square in C. E)

## Discussion

To the best of our knowledge, this is the first article that concurrently quantifies parallel changes in MRI brain features and gut microbiome alterations in rodents self-administering morphine. We present several gut microbiome changes in diversity and taxa along with longitudinal alterations in brain microstructure due to the self-administration of morphine. A subset of animals who underwent cellular analysis finds that rats self-administering morphine show that neuroinflammation could be a pathological mechanism of bidirectional changes in the gut and brain which could be quantified with MRI and complementary analysis of fecal matter. Our work further supports the hypothesis of bidirectional communication between the brain and gut as a mechanism of simultaneous changes due to opioid use. Several studies have looked at the effects of self-administration of opioids such as fentanyl^37,38^, and heroin^39^; however, no studies have examined the impact of the microbiome in addiction paradigms for morphine specifically^14^. In the present study, we show a significant decrease in α diversity measures among rats that self-administer morphine suggesting that the gut-brain bidirectional communication may have a role in morphine use.

The dopamine system is well known for its significant role in the reward circuity associated with substance abuse disorders^40^. Although the dopaminergic pathway is fundamental to comprehend neurobiology of SUDs, recently the gut-brain axis offers a new appreciation into the overarching biological alterations of SUDs^41^. Mice with a depleted microbiome via chronic administration of antibiotics causes impaired cocaine reward processing^42^. Several studies demonstrate the association of opioid use with reductions in the α diversity of the gut microbiota, including this report, indicating an effect of opioids to reduce microbe diversity in the gut. Opioids also alter β diversity showing distinct features of the microbiome composition following use^15^. Subjects receiving addiction treatment show that opioid agonists (i.e., heroin, and prescription opioids) are associated with lower gut microbiome diversity, a finding not observed in those subjects who used agonist + antagonists (i.e., naltrexone), antagonists only, and neither agonists nor antagonists (i.e., other substances not targeting opioids)^42^. Gut microbiome depletion via antibiotics enhanced fentanyl self-administration, while repletion of microbial metabolites via short-chain fatty acid administration reduced fentanyl self-administration in rats^36,37^. Even though several studies establish the relationship between the model of drug use and abuse-altering microbiota, none of those studies have been replicated with morphine.

Morphine upregulates toll-like receptor (TLR) expression levels in small intestinal epithelial cells, sensitizing the small intestine epithelial cells to TLR stimulation ^44^ which induces disruption of tight junctions between epithelial cells increasing gut permeability and bacterial translocation^44^. Animal studies show that morphine treatment results in specific changes to the relative abundance of bacteria^15^. Morphine chronic administration has been shown to reduce diversity of Bifidobacteria and Lactobacillaeae and probiotics enhancing these communities improves behavior and reduced analgesic tolerance to morphine^45^. Morphine administration also causes an increase in the relative abundance of Rikenellaceae,a family from the phylum Bacteroidetes ^46^. Morphine treatment decreases genera *Lactobacillus* and *Clostridium*, which associates with decreased bile salt deconjugation, gut barrier dysfunction, and increased inflammation^43^. Elevated morphine tolerance is associated with abundance of RuminococcaceaeUCG005, and genus. *Ruminococcus 1*^47^. Rats chronically administered morphine display reductions in genus *Bifidobacterium*, during withdrawal and these microbiome alterations have been suggested to be related to inflammatory changes in the amygdala^48^. Morphine administration increases the expression and occupancy of astrocytes in the VTA^49^. The increases in the abundance of *Ruminococcus* sp., and reductions *Lactobacillus* sp., are accompanied with a complimentary activation of microglia^50^. The pro-inflammatory state induced by morphine may be due to an altered Firmicutes-Bacteroidetes ratio leading to gut barrier breakdown and inflammation^51^. In this work we find decreases in Firmicutes and increases in Bifidobacteriaceae, and Erysipelotrichaceae. At the genus level, we observed reduction in *Clostridium*, *Ruminococcus 1*, and *Turicibacter* with an increase in *Bifidobacterium*. A contrast to all other references discussed here is our use of volitional IV self-administration, while all references use either a passive administration or oral consumption of water, or subcutaneous pellets, which makes our result potentially more relevant for a substance use model, and thus could be a more appropriate complement to study brain changes and microbiome alterations due to opioid use disorders.

MRI features correspond to gut composition, and behavior performance in healthy volunteers treated with probiotics in a randomized clinical trial^52^; showing the potential of gut microbiome diversity enhancement in promoting beneficial neural and behavior features. Not only in promoting beneficial features in healthy subjects but also schizophrenia subjects show reduced microbiome diversity mirrored changes in structure and function as measured with MRI^53^. Brain microstructure is usually assessed with DTI (i.e., FA, RD, AD, and MD indices) and is associated with diet-dependent changes in gut microbiota populations, such as *Roseburia* and Barnesiellaceae^54^. Gut microbiota from human subjects with ADHD implanted in mice induces anxiety behavior, and brain microstructure indices of FA and MD in the hippocampus correlate to the abundance of *Eubacterium*^55^. The gut diversity of obese subjects correlates with the FA in the hypothalamus, caudate nucleus, and hippocampus. Also, the abundance of Actinobacteria is associated with DTI indices in the thalamus, hypothalamus, amygdala, and cognitive markers of attention and flexibility^56^. These works highlight the reported relationships between DTI and gut microbiome; however, we did not find a reference that highlight studies of morphine and gut microbiome. Our work identifies several changes in the gut microbiome concurrently with changes in DTI indices, such as a prominent change in the striatum, thalamus, and regions within the frontal lobe.

Neuroinflammation is a possible mechanism linking gut dysbiosis to neural changes due to SUD, implying that complementary gut and brain health markers can be an important tool for assessing neural health. A possible mechanism might be due to drugs of abuse damage to the gut lining, in turn allowing the transfer of bacterial and pathological compounds to induce neuroinflammation ^50,57,58^. Gut disturbances map neuroinflammation signatures to relevant reward circuitry ^59^. Neuroinflammation will induce changes in axonal integrity, and structural connectivity and diffusion imaging studies could provide insights into the potential influence of microglia and neuroinflammatory cascades in neuropsychiatric disorders ^60^. Deficits in intestinal permeability, the gut microbiota may induce inflammation and altered behavior ^61^. Meth exposure promotes microglial activation and cytokine secretion. Meanwhile CCK-8 inhibits Meth-induced microglial activation and IL-6 and TNF-α generation *in vivo* and *in vitro*^62^. Similarly, cocaine alters the gut-barrier composition of the tight junction proteins while also impairing epithelial permeability by upregulating proinflammatory cytokines NF-κB and IL-1β^63^. Repeated administration of meth induces proinflammatory cytokines secretion in the medial prefrontal cortex, striatum, and hippocampus^62^. At the same time, Meth promotes intestinal inflammation by increasing the relative abundance of the pathogenic bacteria in the intestinal tract and reducing intestinal tyrosine hydroxylase protein expression^64^. In our work, we observe several of these brain regions along with a potential link to neuroinflammation in addition to others, such as the brain stem and hypothalamus, not usually reported. Specifically, the brain stem is rich with gut peptide receptors, making them particularly sensitive to signals from the gut^65^.

Our work shows several strengths that make it unique, like the reconciliation of in vivo brain features from MRI to longitudinal gut microbiome analysis linked by confirmation of neuroinflammation in the brain possibly as an important mechanism mediation gut brain axis pathology. Still, we recognize certain limitations of our study. We employ a HARDI (78 diffusion directions) multi-shell sequence (2 b values), which is superior and the ideal method to discriminate anisotropy features in the brain^32,65^. We utilized a relatively low diffusion weighting (b = 500 and 900), which was chosen to accentuate a higher signal-to-noise ratio for all images. Still, future work can employ higher b values to gain higher sensitivity to anisotropic microstructural features. Also, the short access of self-administration (2 hours) affords lower total drug consumption and potentially milder effects than more compulsive elicits methods such as long access or intermittent access. The immunohistochemistry validation can be better estimated with a larger sample size and by acquiring more images with confocal to observe magnified interactions where the high correlation values are on the heat map since the colocalization observed could be blood vessels. In addition, to maintain safe and healthy catheter IV administration, we used cefazolin as a preventative measure, which has been shown to alter microbiome composition^66^. We feel that the overall consistently higher effect in the morphine group and the extensive literature showing neuroinflammation in several brain areas following administrations of drug abuse make us confident of these results.

## Conclusions

With this work, we detected brain and gut changes in vivo and non-invasively in a rodent model of morphine self-administration. Morphine is notoriously known to alter the digestive function and microbiome in humans, as well as in rodent studies using morphine. An understudied aspect of morphine effects in the gut in animal studies is the focus on substance use models, which we attained in this report. In addition, merging MRI with experimental gut techniques, such as 16S rDNA longitudinal sequencing in the same animal, will allow for mapping the trajectory and causal relationships between GBA and SUDs. SUD is one neuropsychiatric disorder that can profoundly gain novel hypotheses and breakthrough insight by leveraging MR and GBA research. Future goals would seek to achieve gut restoration and preserve brain markers of MRI. It is by joining these two in animal studies of SUDs that we may achieve new transcendental insights about opioid use and SUDs that will yield translatable putative biomarkers of drug use that will inform future human studies.

## Acknowledgments

This work was supported by a 2020 Young Investigator Award from the Brain and Behavior Research Foundation. LC was also supported by a grant from the National Institutes of Health 5K25DA047458. We would also like to acknowledge Dr. Bibek Thapa from the HSC Pre-clinical Imaging Core for assistance with the MRI acquisition.

